# Transcriptome-wide investigation of stop codon readthrough in *Saccharomyces cerevisiae*

**DOI:** 10.1101/2020.12.15.422930

**Authors:** Kotchaphorn Mangkalaphiban, Feng He, Robin Ganesan, Chan Wu, Richard Baker, Allan Jacobson

## Abstract

Translation of mRNA into a polypeptide is terminated when the release factor eRF1 recognizes a UAA, UAG, or UGA stop codon in the ribosomal A site and stimulates nascent peptide release. However, stop codon readthrough can occur when a near-cognate tRNA outcompetes eRF1 in decoding the stop codon, resulting in the continuation of the elongation phase of protein synthesis. At the end of a conventional mRNA coding region, readthrough allows translation into the mRNA 3’-UTR. Previous studies with reporter systems have shown that the efficiency of termination or readthrough is modulated by *cis*-acting elements other than stop codon identity, including two nucleotides 5’ of the stop codon, six nucleotides 3’ of the stop codon in the ribosomal mRNA channel, and stem-loop structures in the mRNA 3’-UTR. It is unknown whether these elements are important at a genome-wide level and whether other mRNA features proximal to the stop codon significantly affect termination and readthrough efficiencies *in vivo*. Accordingly, we carried out ribosome profiling analyses of yeast cells expressing wild-type or temperature-sensitive eRF1 and developed bioinformatics strategies to calculate readthrough efficiency, and to identify mRNA and peptide features which influence that efficiency. We found that the stop codon (nt +1 to +3), the nucleotide after it (nt +4), the codon in the P site (nt -3 to -1), and 3’-UTR length are the most influential features in the control of readthrough efficiency, while nts +5 to +9 and mRNA secondary structure in the 3’-UTR had milder effects. Additionally, we found low readthrough genes to have shorter 3’-UTRs compared to high readthrough genes in cells with thermally inactivated eRF1, while this trend was reversed in wild-type cells. Together, our results demonstrated the general roles of known regulatory elements in genome-wide regulation and identified several new mRNA or peptide features affecting the efficiency of translation termination and readthrough.

**AUTHOR SUMMARY:** Of the 64 codons that exist for translation of the genetic code into a polypeptide chain, only UAA, UAG, and UGA signify termination of continued protein synthesis. However, the efficiency of termination is not the same for different mRNAs and is likely influenced by mRNA sequences proximal to these “stop” codons. Here, we sought to expand current understanding of termination efficiency to a genome-wide scale. Our analysis identifies novel mRNA features that may regulate the efficiency of translation termination, including the identities of the stop codon, the penultimate codon, the proximal nucleotides, and the length of mRNA 3’-untranslated region. As ∼11% of human diseases are caused by mutations that introduce premature stop codons, resulting in both accelerated mRNA decay and truncated protein product, our findings are valuable for understanding and developing therapeutics targeting the termination step of mRNA translation.

## INTRODUCTION

Translation of an mRNA into a polypeptide is terminated when the release factor eRF1 interacts with a UAA, UAG, or UGA stop codon in the ribosomal A site and another release factor, eRF3, hydrolyzes GTP and stimulates the polypeptide release activity of eRF1 [1–3]. However, at low frequency, a near-cognate tRNA (nc-tRNA; a tRNA with one stop codon mismatch in its anticodon) outcompetes eRF1 in decoding the stop codon, resulting in the continuation of translation elongation. When the stop codon is located at its normal site at the end of an open reading frame (ORF) elongation will continue into the mRNA 3’-untranslated region (3’-UTR), sometimes producing a C-terminally extended polypeptide product. Such events are termed stop codon readthrough or nonsense suppression [1, 4–6].

The process of readthrough was first characterized primarily in viruses, which utilize this mechanism to increase the protein-coding capacity of their compact genomes [7, 8]. It is only quite recently that C-terminally extended yet functional protein isoforms have been discovered in the proteomes of higher eukaryotes, and their expression is often attributable to readthrough levels that are 10 to 100-fold higher than basal readthrough frequencies [9–18]. In addition, ribosome profiling and phylogenetic investigation of flies, humans, and yeast cells revealed that readthrough is more prevalent than previously anticipated, with efficiencies varying by more than 100-fold for distinct mRNAs [12, 19–22]. These observations indicate that the efficiency of translation termination can be subjected to transcript-specific regulation, and that a key to understanding this regulation may lie in specific mRNA sequences.

Previous studies with reporter systems have shown that the efficiency of termination, and thus the extent of readthrough, is modulated by multiple *cis*-acting elements [4, 5, 23]. The three stop codons themselves differ in termination efficiency, with UGA resulting in the most readthrough, and UAA the least [12, 24]. Nucleotides in the immediate vicinity of the stop codon, including two nucleotides 5’ of the stop codon [25, 26] and up to six nucleotides 3’ of the stop codon in the ribosomal mRNA channel [8, 16, 27–29], have been shown to modulate readthrough efficiency. Additionally, stem-loop structures in the mRNA 3’-UTR are enriched in readthrough-prone transcripts [17, 20, 27, 30]. It remains to be determined whether the effects of these elements are applicable at a genome-wide level for endogenous mRNAs *in vivo*. Transcriptome-wide ribosome profiling studies of flies and yeast have detected readthrough of individual mRNAs [19, 21, 22, 31–33], but detailed dissection of the relationships between readthrough efficiency and *cis*-acting mRNA sequences have only been minimally investigated.

While the stop codon and its immediate 3’ flanking sequences have been studied quite extensively, with structural evidence showing certain nucleotide preferences for optimal interactions with eRF1 motifs and rRNAs [23, 34, 35], other proximal regions have not. Novel readthrough regulatory elements are likely to exist because readthrough has been observed in genes that do not have any of the known readthrough-promoting signals (although there are genes that have readthrough-promoting elements, but do not show detectable readthrough experimentally [10, 15, 20]). Based on studies of other translational control events, such as ribosome stalling [1, 36], several other mRNA features could affect termination and readthrough. The nascent peptide sequences in the ribosomal exit tunnel may affect termination or readthrough efficiency via their interference with the peptidyl transferase center. The proximity of the stop codon to the poly(A)-binding protein (PABP) is also thought to influence readthrough, as PABP is known to interact with eRF3 and enhance termination *in vitro* and *in vivo* [37–39]. These possibilities have yet to be explored with regard to normal termination at a global scale.

Accordingly, we have carried out ribosome profiling of yeast cells expressing wild-type or temperature-sensitive mutant allele of eRF1 and developed bioinformatics strategies to measure readthrough efficiency of individual mRNAs on a genome-wide scale, and to identify mRNA and peptide features which influence that efficiency. Our results demonstrated the general roles of known regulatory elements, such as the identities of the stop codon and the surrounding nucleotides, in genome-wide regulation and identified several new mRNA and peptide features that appear to play a role in translation termination and readthrough, including the penultimate codon in the P site (when a stop codon is in the A site) and the length of the 3’-UTR.

## RESULTS

### Inactivation of eRF1 promotes readthrough of both normal and premature stop codons and accumulation of ribosomes at those codons

We performed ribosome profiling and total RNA sequencing of isogenic wild-type (WT) and eRF1 mutant yeast cells. The latter harbor the *sup45-2* (subsequently referred to as *sup45-ts*) mutation that minimally affects eRF1 function when cells are grown at 25°C, but renders eRF1 inactive at 37°C [40]. Reads between replicates were reproducible, with Pearson’s correlation coefficients (*r*) of 0.96 and 0.99 on average for ribosome profiling and RNA-Seq, respectively (Figure S1A). In addition to the profiling data obtained from these experiments, we also analyzed published ribosome profiling data obtained from eRF1-depleted yeast cells [41] and, for comparison to ribosome occupancies in the 3’-UTR that are not due to termination defects, we included analyses of ribosome profiling data obtained from cells that are depleted of Rli1, a protein involved in post-termination ribosome recycling [42]. Ribosome footprints mapped to genes that have no experimental UTR annotations and genes that overlap by more than 18 base pairs on the same mRNA strand (where read assignment is ambiguous) were discarded from further analyses. With these filters, we were left with 2,693 genes, or ∼41% of the yeast genome, for analyses. This set of genes is referred as the reference gene set.

To explore the consequence of Sup45 inactivation on translation, ribosome footprint reads per kilobase per million (RPKM) that aligned to all regions of each gene of both strains and growth temperatures were plotted against each other. The *sup45-ts* cells grown at 37°C had a higher number of footprints compared to those grown at 25°C or their WT counterpart grown at 37°C (Figure S1B). After normalizing the footprint levels to the mRNA levels to obtain translation efficiency values, *sup45-ts* cells grown at 37°C showed increased ribosome occupancy compared to those grown at 25°C or their WT counterpart grown at 37°C (Figure S1C). These results suggested that the increased footprints were not due to changes in gene expression in response to heat shock, but were likely due to consequences of inactivating eRF1 on translation, including active translation beyond the stop codon.

To see what accounted for the increased ribosome occupancy when eRF1 was inactivated, ribosome footprints were mapped relative to their respective start or stop codons based on their P-site locations and quantified (Figure 1A-D). In all strains, footprints in the coding (CDS) region showed 3-nt periodicity, a profile expected of translating ribosomes. The absence of footprints at the penultimate codon of the open reading frames (ORFs) in *SUP45* cells at 25°C indicated that, with the stop codon in the A site, termination was completed, and ribosomes were dissociated from the mRNAs before they could be captured. The presence of footprints at this position in *sup45-ts* cells at 25°C suggested that the mutation slowed down termination enough for the ribosomes to be captured. When eRF1 was inactivated in *sup45-ts* cells grown at 37°C, or depleted in *sup45-d* cells, a high number of footprints was observed at the penultimate codon, indicating that ribosomes were stalling as they awaited successful termination. Similarly, the high number of footprints in *rli1-d* cells were also consistent with ribosome stalling, as the stop codon had entered the ribosomal A site, but a factor needed for successful recycling was lacking. Additionally, the peaks at approximately 30 nt upstream of the stop codon in *sup45-ts* cells at 37°C and *sup45-d* samples identified ribosomes stacking against those stalled at the stop codon. As a consequence of failed termination or recycling, the mutant libraries, especially from the *sup45-ts* cells at 37°C, had increased footprint reads in the 3’-UTR region compared to those of WT samples (Figure 1A-D, insets), implying stop codon readthrough, frameshifting, or re-initiation events. The mutant samples also showed increased footprint reads at the start codon and decreased footprint reads in the CDS compared to their WT counterparts, observations that are in agreement with the notion that termination and recycling steps are linked to translation initiation [3]. Thus, the overall increase in ribosome occupancy per mRNA in *sup45-ts* cells at 37°C was due to ribosomes stalling at the start and stop codons as well as translation into the 3’-UTR.

**Figure 1.**
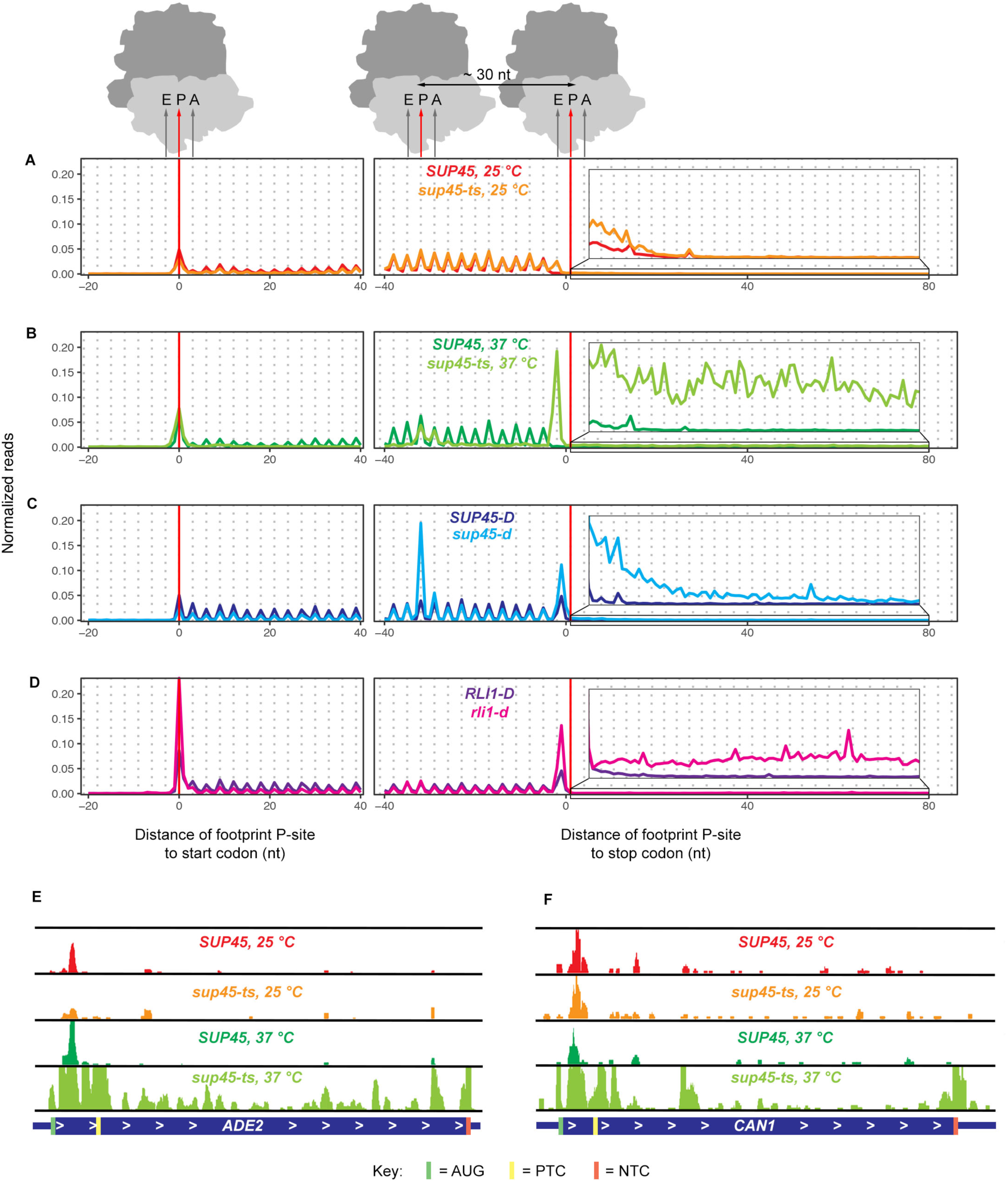
Ribosome occupancy at and beyond the stop codon increases in yeast cells defective in translation termination or ribosome recycling. A-D. Normalized ribosome footprints (2,693 genes with 3’-UTR annotations and without sequence overlapping) aligned at their start or canonical stop codons. Footprints were plotted by the position of their P-sites. Each panel contains the data from WT and its respective mutant. The elevated number of footprints at the penultimate codon of the ORFs in mutant samples demonstrates ribosome stalling when the stop codon was in the A site. (The footprints at this codon and their shift to reading frame +1 in other yeast strains obtained from published data may be attributable to the differences in strain background or sequencing library preparation procedures.) Inset: Magnified view of the region in the box, showing increased 3’-UTR ribosome occupancy in *sup45-ts* cells at 37°C, *sup45-d*, and *rli1-d* cells relative to their respective WT. E-F Read coverage tracks from the Integrative Genomics Viewer (IGV) [67] showing ribosome profiling reads for two PTC-containing genes, *ADE2* (E) and *CAN1* (F), in *SUP45* and *sup45-ts* strains at 25°C and 37°C. Full scale for E equals 50 reads and full scale for F equals 20 reads.

We also examined the effects of Sup45 inactivation on premature translation termination. Two alleles in our strain background, *ade2-1 and can1-100,* contain premature termination codons (PTCs) and their gene specific profiles revealed an accumulation of ribosomes 5’ to the respective PTCs and increased ribosome density in the part of the CDS following the PTCs in *sup45-ts* cells grown at 37°C (compared to *sup45-ts* at 25°C or to WT cells at either temperature) (Figure 1E, F). These observations suggest that readthrough of PTCs also occurs in the absence of functional eRF1. These conclusions were reinforced by analyses of the gene-specific profiles for *RPL28* and *RPS0A* (Figure S2). Intron-containing transcripts from both of these genes enter the cytoplasm and their translation stalls and triggers NMD (nonsense-mediated mRNA decay) when ribosomes encounter PTCs within the respective introns [43, 44]. Figure S2 shows that, for both transcripts, ribosome profiling reads that were generally absent from the respective intron regions in *sup45-ts* cells at 25°C or in WT cells at either temperature were abundant in *sup45-ts* cells grown at 37°C and accumulated mostly at in-frame stop codons (yellow rectangles).

To determine whether the increased ribosome density in regions following the stop codons in eRF1 mutants was due to stop codon readthrough rather than frameshifting or re-initiation events, we calculated the fraction of reads in each of the three reading frames in different mRNA regions: 5’-UTR, CDS, the 3’-UTR region between the canonical stop codon and the first downstream in-frame stop codon (“extension”), and the rest of the 3’- UTR (“distal 3’-UTR”) (Figure 2A). If readthrough occurred, we expected the dominant reading frame in the extension to be the same as in the CDS. Indeed, we saw that frame 0, which was dominant in the CDS, was also dominant in the extension, but not in the distal 3’-UTR of most strains (Figure 2B). When log2 fold-change of footprint fraction in the extension was calculated for each pair of mutant and WT data, *sup45-ts* at 37°C displayed considerable enrichment of frame 0 reads while *sup45-d* and *sup45-ts* at 25°C showed slight enrichment of frame 0 reads relative to their WT counterparts (Figure 2C). On the other hand, *rli1-d* had comparable reading frame preference to its WT counterpart.

**Figure 2.**
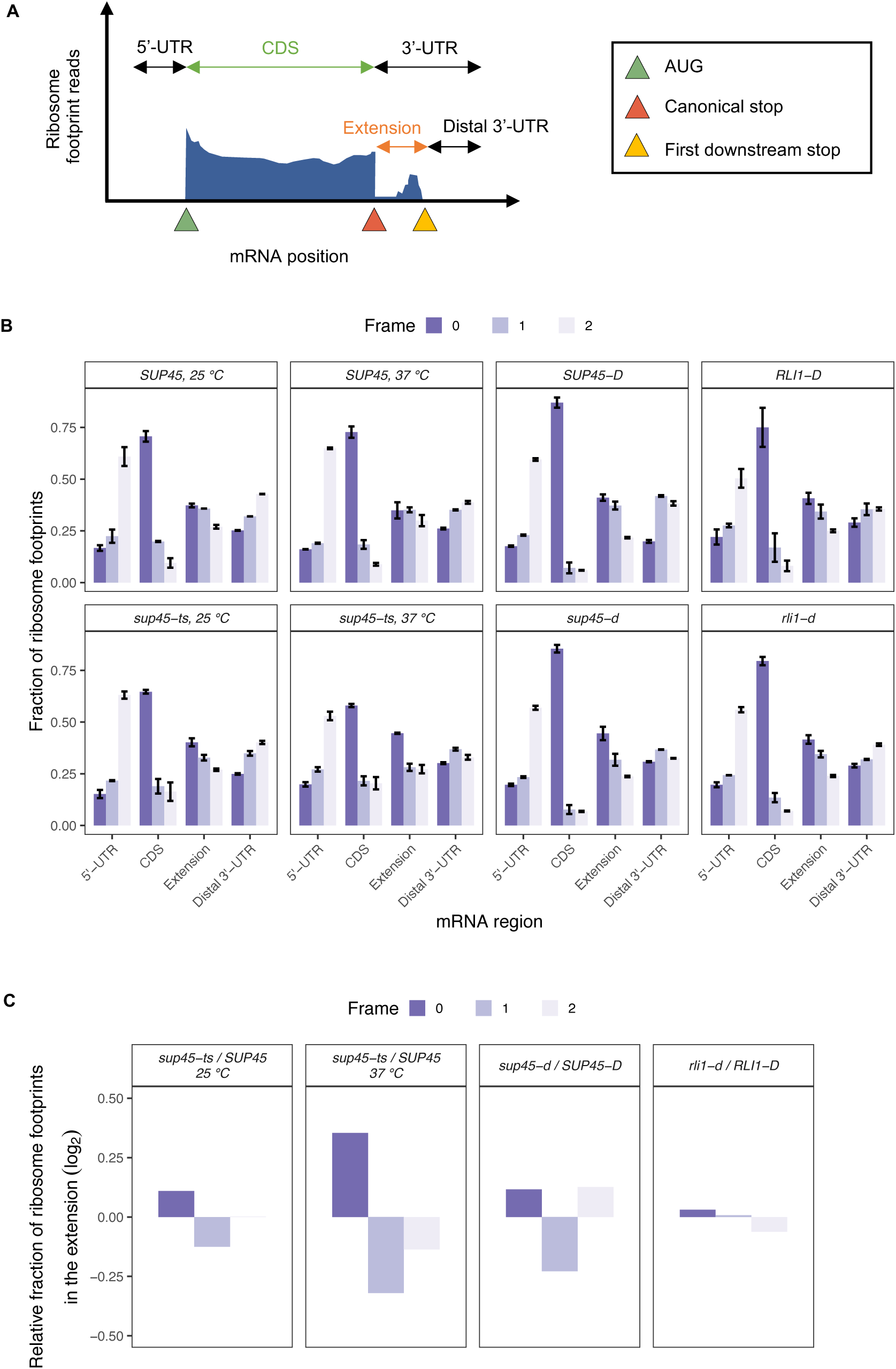
Readthrough events were characterized by maintenance of reading frame when ribosomes bypass the stop codon. **A.** Example of ribosome density plot with labeled mRNA regions of interest (Adapted from Brar and Weissman, 2015, with permission from Springer Nature). **B.** Fractions of reads in each of the three reading frames for different mRNA regions. The results of each sample were the average of 2 or 3 replicates ± SEM. **C.** Ratio of average read fractions (from B) in the extension region of mutant yeast strain over its isogenic WT strain.

In summary, the increased in-frame footprints in the extension region of the 3’-UTR in eRF1 mutants indicated that a lack of functional eRF1 led to stop codon readthrough globally. Hence, we developed formal criteria for the quantitation of readthrough of individual mRNAs and a bioinformatic scheme to assess the genome-wide positive and negative *cis*-acting effects of neighboring sequences to the readthrough of stop codons.

### Analysis pipeline development

To dissect the relationships between readthrough efficiency (Y variable) and mRNA or nascent peptide features (X variables), we used the random forest machine learning approach [45, 46] to first narrow down the candidate features. We defined each of the variables as follows:

#### Defining readthrough efficiency (Y variable)

Readthrough efficiency in the context of ribosome profiling can be loosely defined as the ratio of ribosome density (footprint count normalized to the length of the region) in the 3’-UTR to ribosome density in the CDS; however, the nature of the mutants should be taken into account to ensure accuracy and minimize noise. First, due to ribosome stalling and queuing at the start and stop codons in mutant strains (Figure 1A), ribosome density in the CDS may be overestimated; therefore, we excluded footprints whose P-sites fall into the first 15 and the last 33 nucleotides of the CDS of each gene in all samples. Second, as readthrough is by definition in-frame with the CDS (Figure 2B), only footprints that were in frame 0 were considered. The adjusted readthrough efficiency calculation, log-transformed, is shown in Figure 3A.

**Figure 3.**
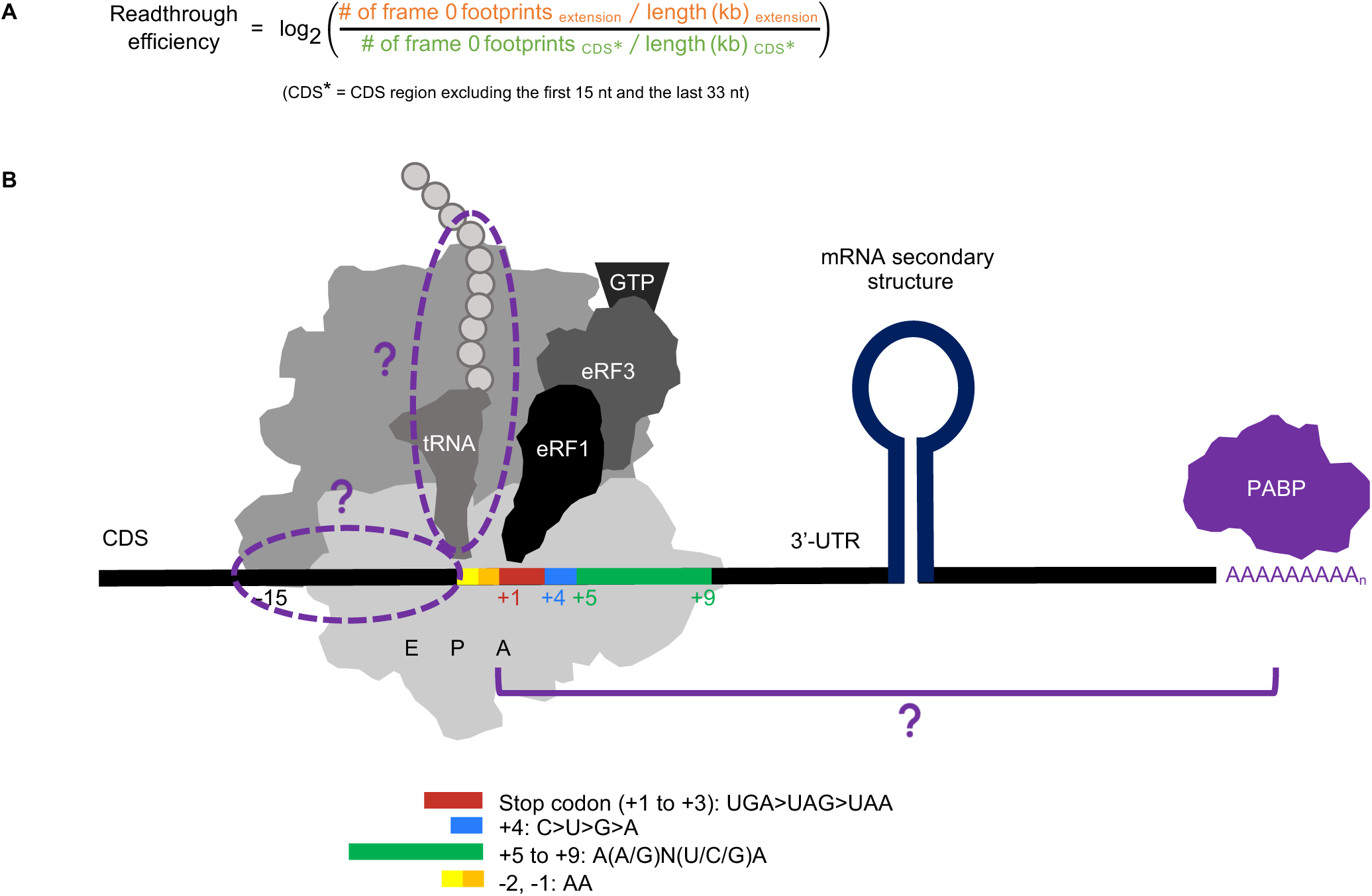
Definition of X and Y Variables. **A.** Readthrough efficiency calculation formula, which takes into account the ribosome pile-up at NTCs and reading frame preference. **B.** mRNA and nascent peptide features of interest in the context of the ribosome and related translation termination components. Previously identified variables are colored and labeled. Hypothesized variables are in purple and marked with “?”. Details of how each variable is measured are in Table 1. (Figure adapted from Rodnina et al., 2019, with permission from Oxford University Press).

**Table 1.**
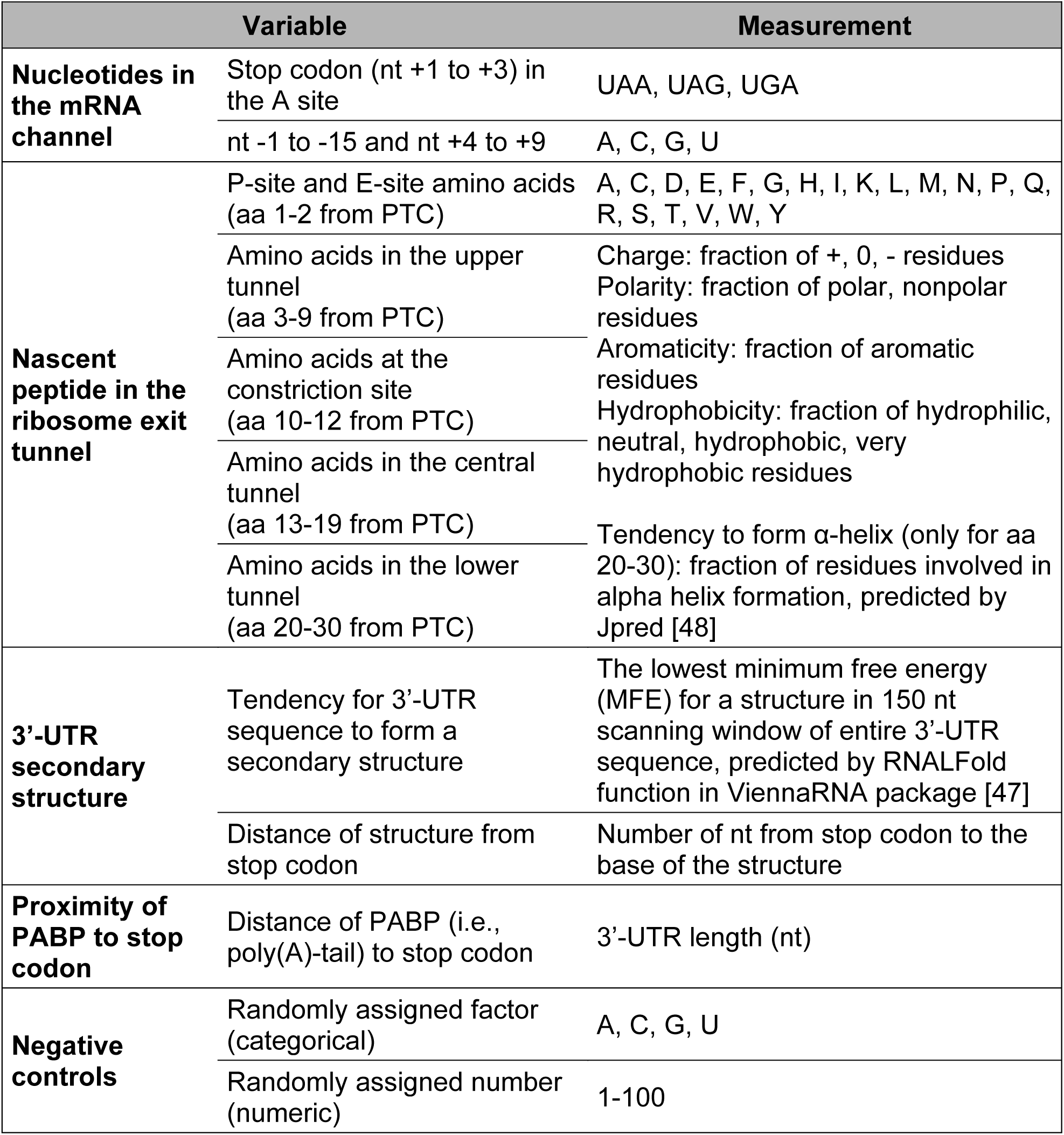
Assessed X variables and their modes of measurement.

Additional sources of noise in the data include: 1) poorly-expressed genes that result in an exaggeration of the readthrough efficiency ratios and 2) genes lacking footprints in the 3’-UTR region, for which it was challenging to distinguish lack of readthrough from insufficient sequencing depth. Inclusion of these genes confounded previous analyses and led to conflicting results. For instance, genes belonging in the “leaky” set (genes that had 3’-UTR reads, i.e. readthrough) were highly expressed compared to genes in the “non-leaky set” (genes lacking 3’-UTR reads), but within the “leaky” set, readthrough rate was negatively correlated with gene expression [22]. Therefore, we discarded genes with both RPKM of CDS < 5 and RPKM of 3’-UTR < 0.5 [42] from further analyses.

#### Defining mRNA and nascent peptide features (X variables)

To date, mRNA features affecting stop codon readthrough have frequently been identified 3’ of the stop codon, but we also considered the possibility that mRNA sequence on the 5’ side of the stop codon and the nascent peptide in the exit tunnel may contribute to variation in readthrough efficiencies (Figure 3B). These features can either be directly documented or inferred from the mRNA sequences (Table 1). Therefore, we used established algorithms to predict RNA secondary structure [47] and nascent peptide α- helix formation [48].

Other than the mRNA features listed in Table 1, many characteristics of the mRNAs (such as expression levels, codon optimality, translation efficiency, and gene length) were previously correlated with readthrough efficiency measured by ribosome profiling [21, 22, 32]. We performed similar analyses involving pairwise correlation between readthrough efficiency, gene expression level (determined by RSEM of RNA-Seq data), translation efficiency (ribosome density in the CDS divided by gene expression level), length of transcript and mRNA regions, and codon optimality (tRNA adaptation index [49, 50]) (Figure S3). We found that, except for *sup45-ts* at 37°C, readthrough efficiency was negatively correlated with gene expression and codon optimality, consistent with previous results [21, 32]. This correlation may have an evolutionary, rather than a mechanistic, explanation; for example, the inverse relationship between gene expression and readthrough efficiency may result from more deleterious effects of readthrough from highly expressed genes than from poorly expressed genes, and thus highly expressed genes evolved to have lower readthrough efficiency [32]. Since gene expression level was positively correlated with codon optimality across the ORF, where mRNAs with optimal codons are more stable and hence more abundant than those with non-optimal codons [51], we could not make a distinction whether the observed negative correlation between readthrough efficiency and codon optimality was simply due to increased mRNA abundance or was instead mechanistically relevant to the readthrough process. We therefore did not include gene expression and codon optimality in further analyses. Moreover, readthrough measurements in general should be interpreted with caution and due consideration given the caveats described above.

### Random forest analyses reveal stop codon identity, P-site amino acid identity, and 3’-UTR length as important factors in predicting readthrough efficiency

In order to identify without bias the mRNA and nascent peptide features affecting readthrough efficiency, we used two independent approaches involving the random forest algorithm [45, 46]: 1) random forest regression to predict readthrough efficiency based on the X variables, and 2) random forest classification to differentiate between “High” (top 15%) and “Low” (bottom 15%) readthrough genes based on the X variables. The number of genes in each type of analysis can be found in Table S4. A regression model and a classification model with 5-fold cross-validation were created for each sample using the same tuning parameters. For the regression model, we used Normalized Root Mean Square Error (NRMSE) as a measurement of predictive ability of X variables. We observed NRMSE of approximately 0.14 across all samples, indicating that the average difference between predicted and actual readthrough efficiency is 14% (Figure 4A). For the classification model, Area Under the Receiver Operating Characteristics (AUROC) was used as a measurement of model capability at distinguishing between the “High” and “Low” groups. AUROC of approximately 0.7 was observed across all samples, therefore the models had a 70% chance of classifying genes into the correct group (Figure 4B).

**Figure 4.**
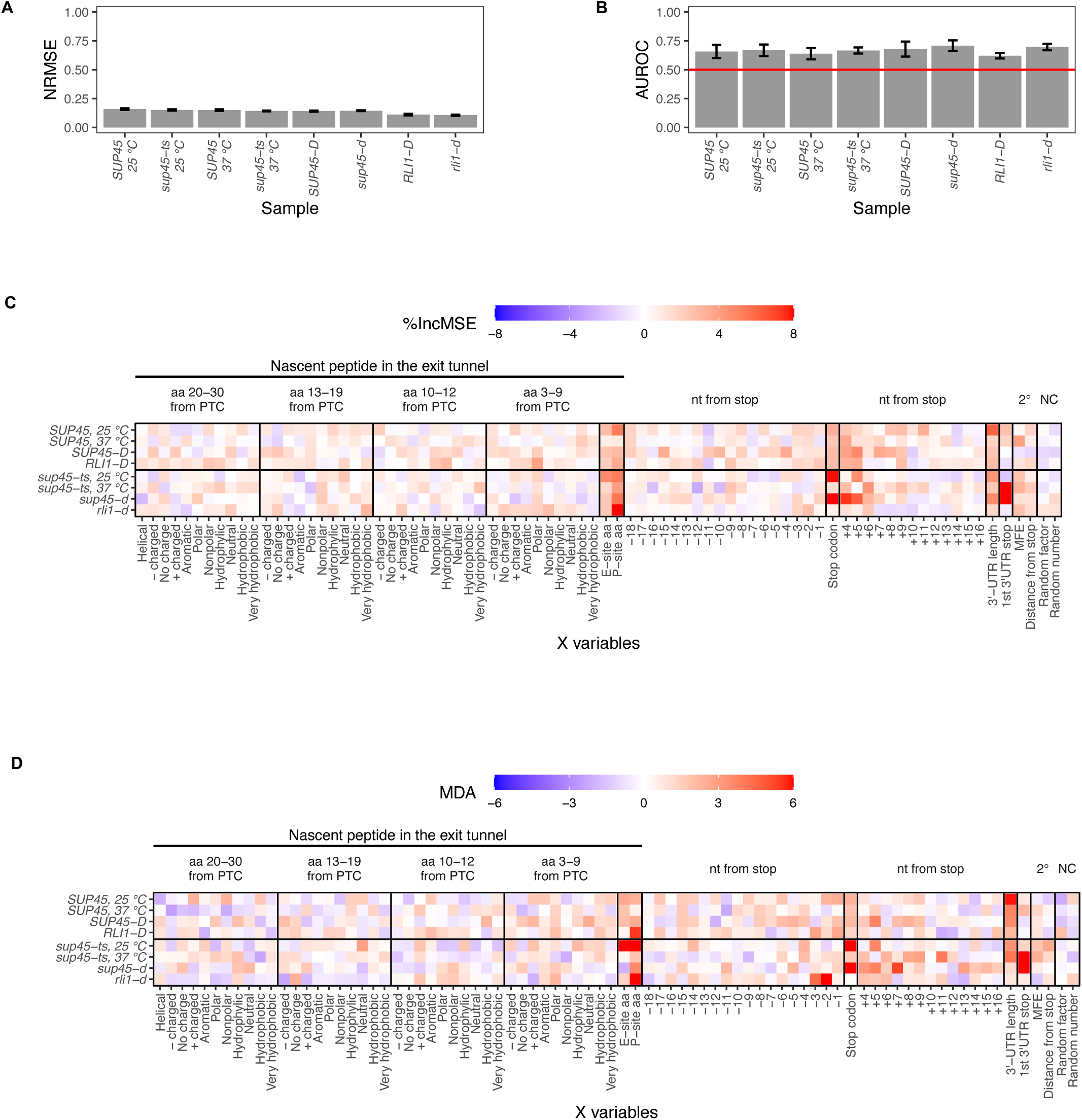
Random forest identified stop codon identity, P-site amino acid identity, and 3’-UTR length as factors critical in readthrough efficiency prediction. **A.** Average Normalized Root Mean Square Error (NRMSE), measurement of model accuracy for random forest regression, across 5-fold cross-validation. **B.** Average Area Under the Receiving Operator Curve (AUROC), a measurement of model accuracy for random forest classification, across 5-fold cross-validation. The value of 0.5 (red line) means the classification by the input X variables is no better than random chance. **C.** and **D.** Feature importance extracted from the random forest models. The relative importance of a feature is represented by percent increase in mean square error (%IncMSE) for regression (C) and Mean Decrease Accuracy (MDA) for classification (D), which is percent increase in error that results from permuting the feature. The higher the %IncMSE or MDA is, the more crucial the feature is in predicting readthrough efficiency or correctly classifying genes into groups. Arbitrary continuous and discrete values randomly assigned to the genes (random number and random factor) are used as negative controls.

To determine which of the X variables were most responsible for the predictions, we permuted each X variable for the regression and classification models and assessed its importance by calculating either percent increase in mean square error (%IncMSE) or Mean Decrease Accuracy (MDA). %IncMSE is the calculation of percent increase in prediction error, while MDA is the percent increase in misclassification, when the X variable was permuted. Thus, the higher the %IncMSE or MDA value is for an X variable, the more critical that variable is in predicting readthrough efficiency. As baselines (negative controls) for meaningful %IncMSE and MDA values, a number between 1-100 (random number) and one of four categorical values (random factor) were assigned to each gene before model training. Because the assignment was random, these two variables should have no influence on the prediction. Indeed, random number and random factor were among the X variables with %IncMSE and MDA values close to zero (Figure 4C, D). Reassuringly, in both the regression and classification approaches, the variables previously known to affect readthrough efficiency, such as the identities of a stop codon, nucleotide (nt) +4 and +5 were among the variables with %IncMSE and MDA values above the baseline in most samples, arguing that our pipeline was capable of distinguishing relevant mRNA features. In addition to the stop codon identities, 3’-UTR length and the P-site amino acid (or tRNA or codon) also had one of the highest %IncMSE and MDA values in most samples. The nt -3, and -2 in the P-site only had high MDA values in the ribosome recycling mutant, *rli1-d,* suggesting that they may play a role in recycling; however, this is only true in the classification model.

Interestingly, the first downstream stop codon in the 3’-UTR (“1st 3’-UTR stop” in Figure 4C, D) had high %IncMSE and MDA values only in *sup45-ts* at 37°C and *sup45-d* samples. This observation is due to ribosome pile-up at those stop codons, as with canonical stop codons (Figure 1B, C). The identity of this stop codon affects read count in the extension region – its high %IncMSE and MDA value reinforces the role of stop codon identity in readthrough but not mechanistic relevance to readthrough occurring specifically at the canonical stop codon.

In summary, random forest analyses reveal identities of the stop codon, P-site amino acid (or tRNA or nucleotides), and 3’-UTR length as mRNA features that control genome-wide readthrough efficiency.

### Readthrough-promoting stop codons and nucleotides at positions +4 to +9 occur at a genome-wide level

Previous studies using synthetic dual reporters have shown that the UGA stop codon and certain nucleotides at positions -2, -1, +4 to +9 are the most permissive for stop codon readthrough (Figure 3B) [5, 8, 16, 23–29]. Since the random forest analyses also identified the stop codon identity and some of these nucleotides as features that, when permuted, led to higher prediction error than baseline, we wondered whether these same stop codon and nucleotide identities were also readthrough-permissive or readthrough-inhibiting at a genome-wide level. To answer this question, the genes were divided into 3 groups for analysis of stop codon identity or 4 groups for analyses of individual nucleotides at each position and Wilcoxon’s rank sum test was used to compare the median of genes in each group to the overall sample median. The differences between group median and sample median were shown as a heatmap (Figure 5A). The significance values from Wilcoxon’s rank sum test were represented as the size of the tiles, where a larger tile depicted p-value < 0.05. As expected, the distribution of random factor variables was not significantly different between the groups and sample median and showed no particular pattern of higher or lower readthrough efficiency across all samples.

**Figure 5.**
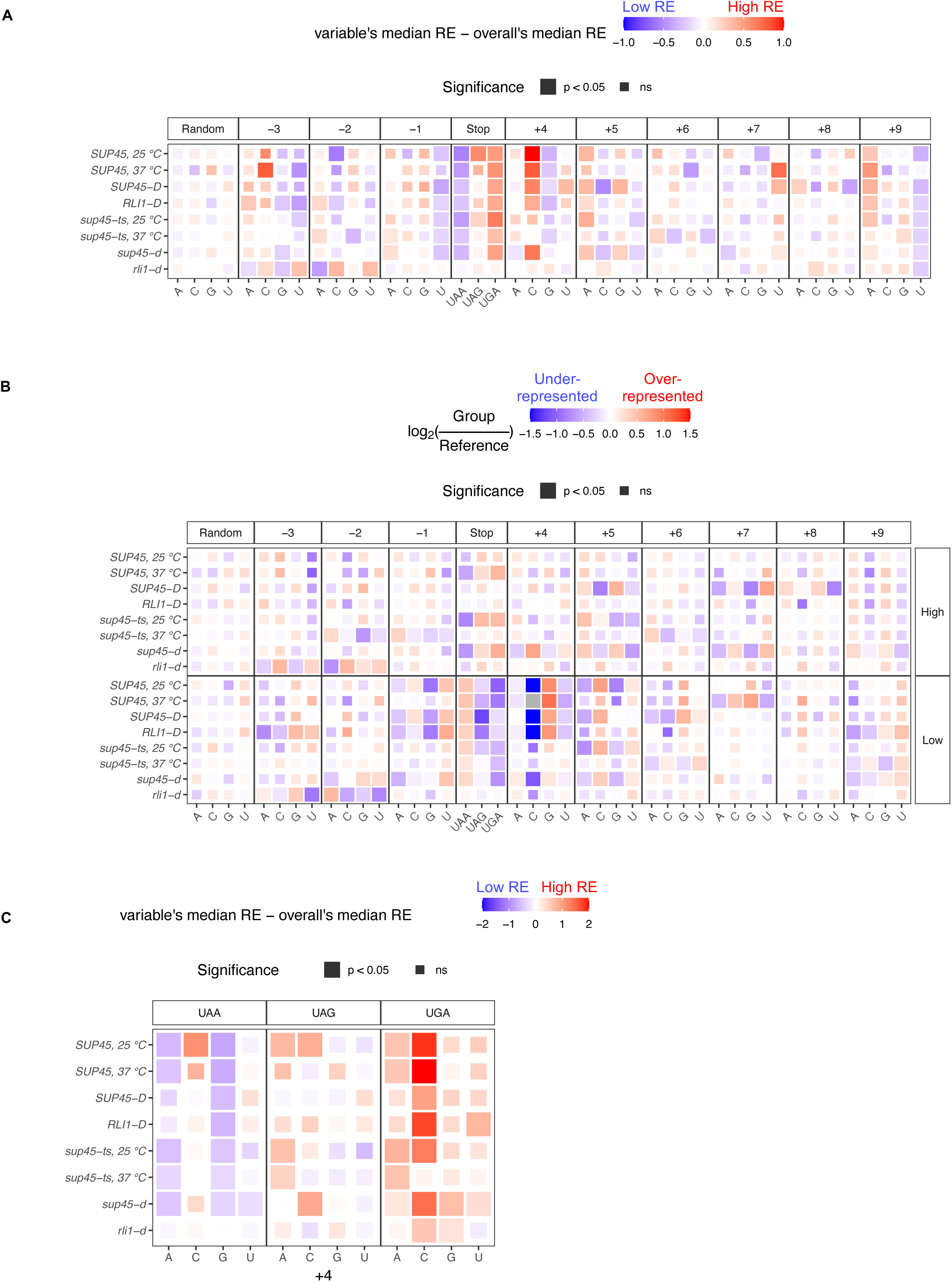
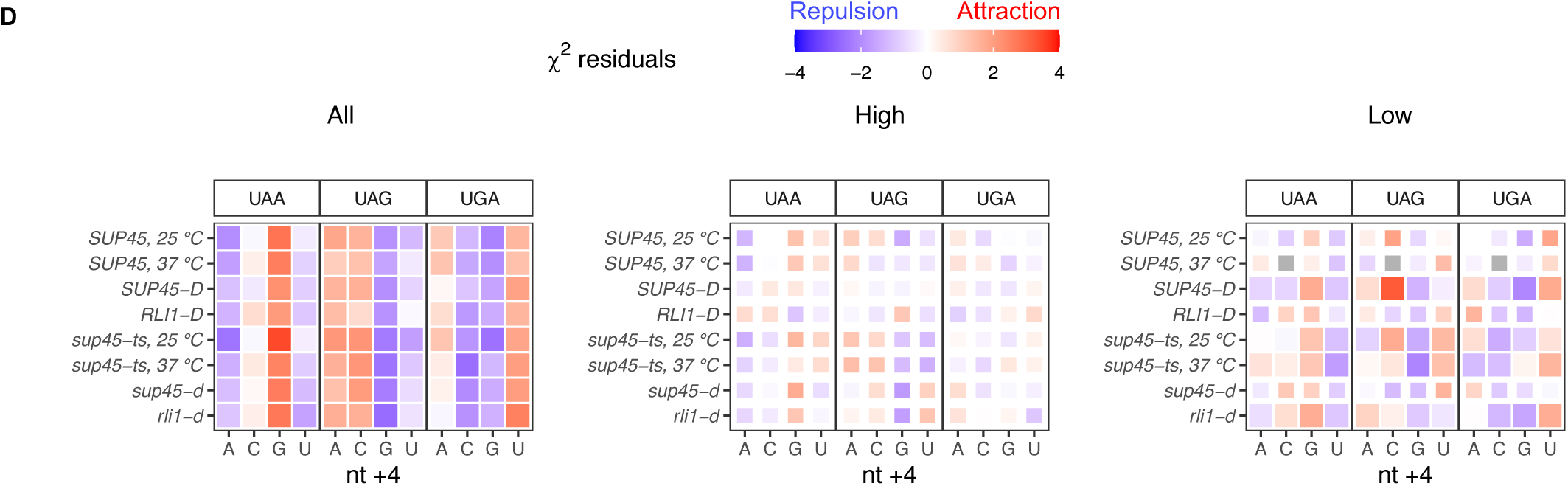
Known effects of stop codon and surrounding nucleotide identities on readthrough efficiency occur at a genome-wide level. **A.** Heatmap of median readthrough efficiency of genes containing particular stop codon or nucleotide relative to median readthrough efficiency of all genes in the sample. Positive value (red) indicates that the group of genes had higher readthrough efficiencies compared to the sample median, while negative value (blue) indicates lower readthrough efficiencies. Wilcoxon’s rank sum test was used to determine whether the difference in group and sample media was significant. Significant difference was represented as a larger tile. **B.** Heatmap demonstrating over-representation (red) or under-representation (blue) of a stop codon or nucleotide in “High” or “Low” readthrough groups relative to the frequency in the reference gene set. χ^2^ goodness of fit test with Bonferroni correction was performed to compare usage distribution between the “High” readthrough group, “Low” readthrough group, and the reference. Significant difference was represented as a larger tile. Grey tiles signify that there was zero observation of that nucleotide (essentially under-represented), and log2 calculation and statistical analysis could not be performed. **C.** Heatmap of median readthrough efficiency of genes containing particular combination of stop codon and nt +4 relative to median readthrough efficiency of all genes in the sample. Statistical analysis as in A. **D.** Heatmap showing residuals of χ^2^ test of independence between stop codon and nt +4 identities. Significant values (p < 0.05), represented by larger tile size, indicate the significant association between stop codon and nt +4 identities. Positive (red) residuals specify positive association (attraction) and negative (blue) residuals negative association (repulsion) between variables.

The variables that showed the strongest pattern and significance were the stop codon and the nucleotide immediately downstream (nt +4). Consistent with the notion that UGA is the most readthrough-permissive stop codon and UAA is the strongest terminator [24], genes with UAA as stop codons had significantly lower readthrough efficiencies while genes with UGA had significantly higher readthrough efficiencies compared to the sample median in all samples except for *rli1-d* cells. For nt +4, genes with C at this position had significantly higher readthrough efficiencies, and those with G had slightly lower readthrough efficiencies. These observations are in line with previous data where C was reported to allow the most readthrough and G the least [5, 24, 27, 28].

For nucleotides further into the mRNA channel 3’ of the stop codon (nt +5 to +9), only nt +9 showed clear trends: genes with A had higher readthrough efficiencies (i.e., readthrough-permissive), consistent with previous reports, and genes with U had lower readthrough efficiencies (i.e., readthrough-inhibiting). At position +5, although the differences between groups’ readthrough efficiencies and overall readthrough efficiencies were not significant in every sample, genes containing adenine had a uniform pattern of higher readthrough efficiency compared to sample median (unlike random factor), so we conclude that A at nt +5 was relatively readthrough-permissive.

Adenine at nt -1 and -2 has been shown to be readthrough-permissive [25, 26], but this was not reflected in our current analysis (Figure 5A). Instead, we observed that U at nt -1 was readthrough-inhibiting, and A and C but not G and U at nt -3 were readthrough-permissive. It is noteworthy that nt -3 was identified as having a significant role along with nt -2; this may mean that the amino acid they encode or its tRNA, rather than the nucleotides themselves, mechanistically affect the termination process. Thus, we also explored these three nucleotides as a codon (see below).

It is notable that most of the patterns for nt -3 and -2 in *rli1-d* sample did not match with other samples. In fact, some of them appeared to be the opposite. Since other samples, even the *sup45* mutants, had wild-type Rli1, the opposite patterns in *rli1-d* sample may suggest that nt -3 and -2 (or the amino acid they encode) played a role in a ribosome recycling step rather than the termination step. This also influenced the number of ribosomes translated into the 3’-UTR in *rli1-d*, as shown in the feature importance plot (Figure 4D).

To see whether we could extract more comprehensive information on these nucleotides, we used a different approach involving only the “High” and “Low” readthrough genes used for the classification models. χ^2^ goodness of fit tests with Bonferroni correction were used to compare the frequencies of UGA, UAG, UGA, A, C, G, and T at each nucleotide position between the “High” readthrough group, “Low” readthrough group, and the reference gene set (2,693 genes). For data presentation purposes, the frequencies were converted into fractions and log2 ratios of “High” or “Low” groups to the reference were computed, as shown in the heatmap (Figure 5B). A positive log2 ratio (red) indicates an over-representation while a negative log2 ratio (blue) indicates an under-representation of X in the group compared to reference. The patterns, especially those of the “Low” readthrough group, corroborated our previous approach (Figure 5A). For instance, genes with C at the nt +4 position had higher readthrough efficiencies than the sample median (Figure 5A), and within the “Low” readthrough group (Figure 5B), C was severely under-represented (blue tile) or even absent (grey tile). Moreover, we were able to rank the nucleotides from readthrough-permissive to readthrough-inhibiting using this approach. Although the patterns of either groups were not significantly different from reference based on the χ^2^ analysis, uniform patterns could be observed across all samples for most nucleotide positions (compared to random factor). For nt +5, the pattern allowed us to extrapolate that following readthrough-permissive G, were increasingly readhthrough-inhibiting A, U, and C (which was most associated with termination). This information was difficult to obtain in the dual reporter assays because readthrough events were so low that they were indistinguishable from each other. Information compiled for other nucleotide positions is in Table 2.

**Table 2.** Comparison of nt -3 to +9 results (based on Figure 5) to previous data (based on review by Rodnina et al., 2019). The order of nucleotide/stop codon is from most to (“>”) least permissive to readthrough.

| Position | Literature<br>(Rodnina et al., 2019) | This study |
| --- | --- | --- |
| -3 | - | A/C > G > U |
| -2 | A | A > G > U > C |
| -1 | A | A > G > C > U |
| Stop codon (+1, +2, +3) | UGA > UAG > UAA | UGA > UAG > UGA |
| +4 | C > U > G > A | C > U > A > G |
| +5 | A | A > G > U > C |
| +6 | A/G | A/C > G/U |
| +7 | N | U |
| +8 | U/C/G | G > A/U > C |
| +9 | A | A > C/G > U |

Next, we asked how particular combinations of stop codon and nt +4 identities, the two strongest variables among all the nucleotides, affect readthrough efficiency together. Genes were divided into groups based on their stop codon and nt +4 identities and the same analysis as described in Figure 5A was performed. We found that genes containing UGA, the most readthrough-permissive stop codon, followed by C, the most readthrough-permissive nt +4, had the highest readthrough efficiency relative to the overall median. UGA followed by G, the least readthrough-permissive nt +4, reduced readthrough efficiency but not to the level of overall median (Figure 5C). On the other hand, genes containing the most readthrough-inhibiting combination, UAAG, had the lowest readthrough efficiencies while UAAC brought readthrough up to a level that was equal to that of the overall median.

To test whether particular stop codon and nt +4 identities tend to occur together at different readthrough rates, a χ^2^ test of independence was performed using all genes (“All”), genes in the “High” readthrough group, or genes in the “Low” readthrough group. We found that in general (“All”), the association between stop codon and nt +4 identities was significant in all samples. Readthrough-inhibiting features, UAA and G, appeared together more frequently than the expected frequency (positive residuals) while readthrough-promoting features, UGA and C, tended to repel each other (negative residuals) (Figure 5D, left panel). Similar but weaker associations were seen among genes with “Low” readthrough, but no association was observed among those with “High” readthrough. From an evolutionary standpoint, this analysis supported a previous proposal stating that genes have evolved to minimize stop codon readthrough [32], as low readthrough features tended to coexist but high readthrough features tended to exclude each other.

To summarize, we showed that readthrough-promoting stop codon and nucleotides at positions +4 to +9 previously determined in reporter assays also occur at a genome-wide level, with some combinations of stop codon and nt +4 identities selected for or against each other. Additionally, we identified readthrough-inhibiting nucleotides refractory to previous studies due to the detection limits of reporter assays.

### Specific combinations of nucleotides and tRNA in the ribosomal P site may influence readthrough efficiency

Random forest identified the P-site amino acid to be a key factor in readthrough efficiency prediction, and we observed patterns of nucleotide usages at position -3, -2, and -1; however, it was unclear whether termination was influenced by the nucleotide themselves, the amino acid they encode, the tRNA that decoded them, or a combination of these three possibilities. Therefore, we performed analysis as described in Figure 5A, but as codons instead of individual nucleotide positions (Figure 6A) and were able to identify codons associated with genes having significant increases or decreases in readthrough efficiencies compared to the sample median.

**Figure 6.**
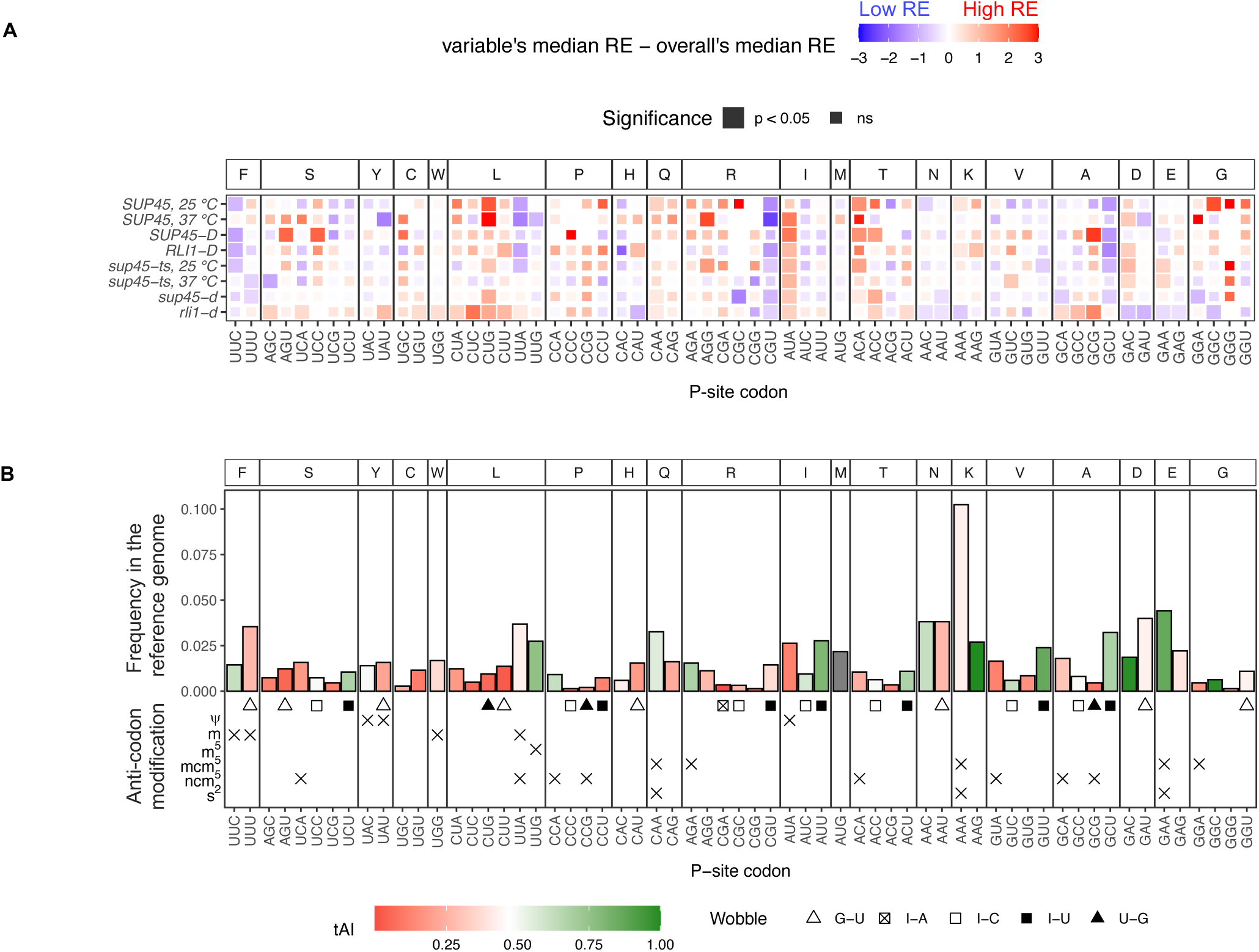
Specific codons in the P site were associated with higher or lower readthrough efficiencies. **A.** Heatmap of median readthrough efficiency of genes containing a particular triplet codon in the ribosomal P site. Positive value (red) indicates that the group of genes had higher readthrough efficiencies compared to sample median, while negative value (blue) indicates lower readthrough efficiencies. Wilcoxon’s rank sum test was used to determine whether the difference in group and sample media was significant. Significant difference was represented as a larger tile. **B.** General information regarding the codons. The Y-axis shows frequency of codon usage in the P site among the reference gene set. Color of the bar represents tRNA adaptation index (tAI), a measurement for codon optimality where 0 is non-optimal and 1 is optimal [49, 50]. Below the bar plots are wobble pair information and anti-codon modifications.

From this analysis, two particular nucleotide combinations were likely responsible for low readthrough efficiencies, namely the C/G and U combinations and the UU and A/C combinations. In the first group, CGU and GCU were codons that appeared in genes with significantly lower readthrough efficiencies in most samples even though they code for different amino acids and were clearly distinct from their respective synonymous codons. Remarkably, genes containing CUG (leucine), which had exactly the same nucleotide composition, showed higher readthrough efficiencies compared to sample median. These opposite trends suggested that not only specific combinations of nucleotides, but also specific positional arrangements have to be optimal for efficient termination. In the second group genes containing UUA and UUC had lower readthrough efficiencies but only in WT conditions (WT strains and *sup45-ts* at 25°C). Similar to the C/GU case, UUA and UUC had the same pattern despite coding for different amino acids and were distinct from their respective synonymous codons.

In both cases, the amino acid identity was ruled out as a possible mechanism. We wondered whether these two groups of codons share certain properties that may explain their occurrence in low readthrough genes and explored P-site codon usage frequency in the genome, tRNA adaptation index (tAI) [49, 50], tRNA anti-codon modifications [52], and wobble pairs (Figure 6B). None of these properties were shared exclusively among the four codons, suggesting the mechanisms by which UUA/UUC and CGU/GCU influence readthrough efficiency were different. Nevertheless, the role of tRNA properties could not be ruled out. It is likely that the nucleotides and tRNAs together interact with the translation termination components in such a way that is optimal for a termination event to occur.

At the other end of the spectrum, codons that appeared in genes with significantly higher readthrough efficiencies include AUA, ACA, ACC, CUG, and GAC. Arguments similar to those made for the low readthrough set of codons applied here. First, the amino acid identity could be ruled out. Although ACA and ACC encode the same amino acid (threonine), it could be ruled out as a possible mechanism because other threonine codons showed different patterns. All five codons did not share any tRNA properties in common; however, individualized nucleotide-tRNA property combinations could not be ruled out as possible mechanisms.

As with nt -3 and -2, many P-site codons (e.g., GAC, CUA, CUC, UUA) in the *rli1-d* mutant had either the opposite effect on readthrough efficiencies or opposite level of significance compared to the eRF1 mutants and WT cells. Thus, it is possible that these codons affect the recycling step rather than termination step.

In summary, we identified four codons (CGU, GCU, UUA, and UUC) associated with lower readthrough efficiencies and five codons (AUA, ACA, ACC, CUG, and GAC) associated with higher readthrough efficiencies. It is unlikely that amino acid identity by itself is the reason for this observation; rather, specific combinations of nucleotide and tRNA properties likely influence termination and readthrough.

### Readthrough efficiency increased with 3’-UTR lengths in the eRF1 mutants, but not in wild-type or recycling factor mutant cells

Poly(A)-binding protein (PABP, Pab1 in yeast) has been shown previously to interact with eRF3, enhancing termination efficiency *in vitro* [37]. Consistent with these observations, deletion of the *PAB1* gene in yeast, or extension of the mRNA 3’-UTR length downstream of a PTC, leads to inefficient termination and readthrough *in vivo* [39]. Accordingly, we asked whether readthrough efficiency and 3’-UTR length are positively correlated. A weak but significant positive correlation between readthrough efficiency and 3’-UTR length was observed in *sup45-ts* cells at 37°C and *sup45-d* cells while a negative correlation was observed in the respective WT strains (Figure 7). These results hinted that Pab1’s expected role in aiding termination became prominent only when eRF1 activity was inefficient, a possibility consistent with recent studies of context dependent translation termination in Ciliates [38].

**Figure 7.**
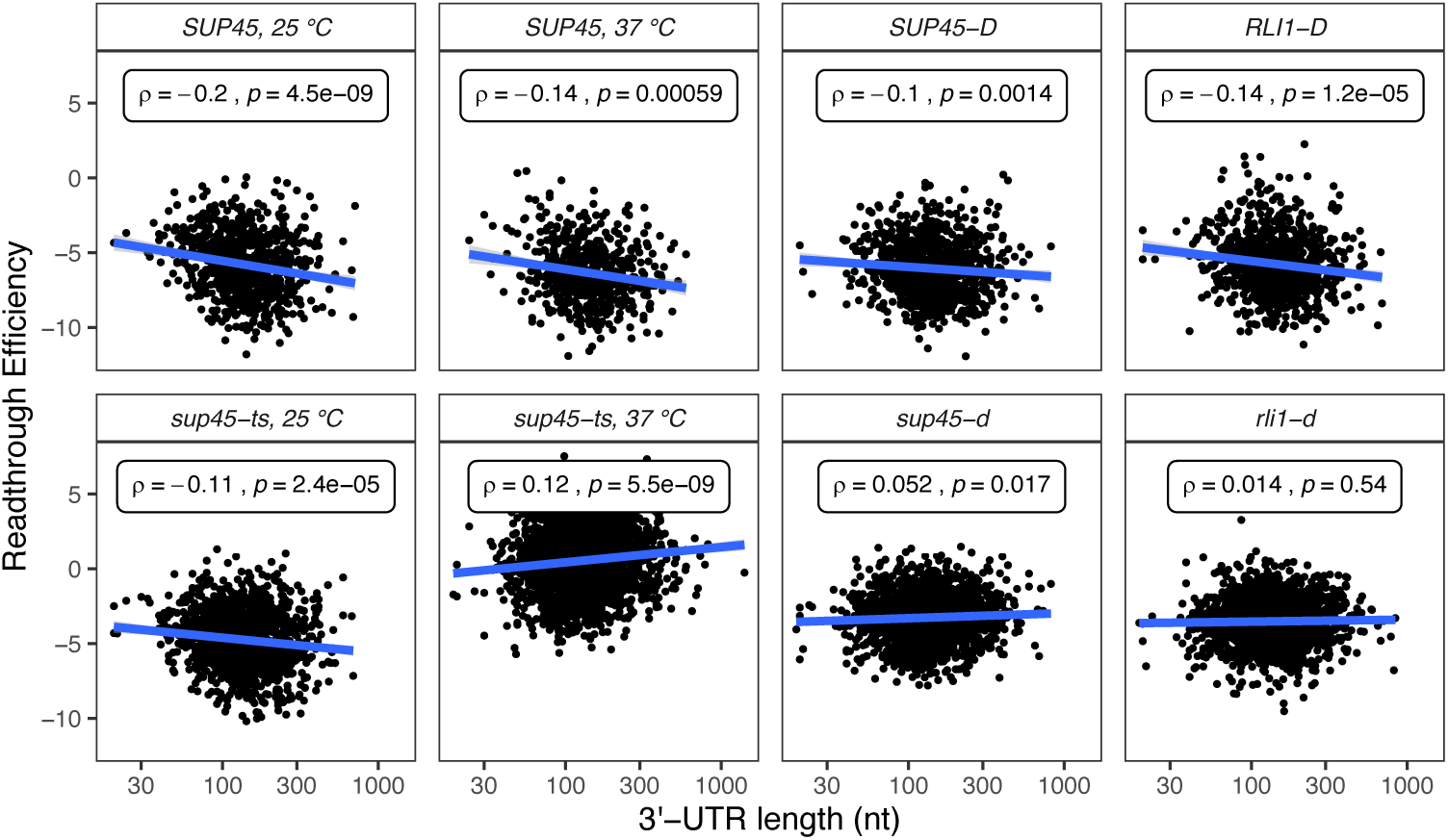
Readthrough efficiency increased with 3’-UTR lengths in eRF1 mutants, but not in wild-type or recycling factor mutant cells. Scatter plots of readthrough efficiency vs. 3’-UTR length. Spearman’s correlation coefficient (ρ) and p-value were calculated for each sample.

## DISCUSSION

In this study, we analyzed ribosome profiling data of WT and mutant yeast strains defective in termination or ribosome recycling to identify mRNA or peptide features that influence readthrough efficiency at a genome-wide level. First, we characterized the phenotypes of eRF1 mutant cells, which included ribosomes stalling at start and stop codons and increased in-frame footprints in 3’-UTRs compared to wild-type cells (Figures 1, 2B, C). We took these phenotypes into account when calculating readthrough efficiency of each transcript (Figure 3A) and excluded any transcripts that lack 3’-UTR annotation or sufficient footprints in the 3’-UTR from further analyses. Although these considerations left us with a smaller number of analyzable data points, we were able to reduce noise in the data that confounded a previous analysis attempt [21, 22]. With our analysis strategies, we demonstrated that the readthrough-promoting stop codon and proximal nucleotides previously determined in reporter assays [16, 27–29] are not reporter-specific, but also occur at a genome-wide level in yeast for endogenous mRNAs (Figure 5). Moreover, our analyses identify readthrough-inhibiting nucleotides that were refractory to previous studies using reporter gene assays because the low level readthrough was below the detection threshold.

Two novel mRNA features that we found to be major determinants of readthrough efficiency were the codon in the ribosomal P site (when a stop codon is in the A site) and 3’-UTR length. For the P-site codon, our results did not support previous conclusions of AA being the most readthrough-permissive nucleotides at position -2 and -1 [5, 25, 26] when we analyzed the data by nucleotide position (Figure 5, Table 2). Moreover, when we analyzed nt -3, -2, and -1 together as a codon in the P site, we did not find codons NAA to be significantly associated with higher readthrough. Instead, we found that genes with AUA, ACA, ACC, CUG, and GAC as their penultimate codon had higher readthrough efficiencies compared to the sample median (Figure 6A), suggesting that there may be properties in the P site other than adenines that could influence readthrough. Previous studies ruled out properties of the last amino acid residue (encoded by the P-site codon) as features affecting readthrough, but reported conflicting results for tRNA in the P site [25, 26]. We were also able to rule out amino acid properties, and although we could not pinpoint the importance of tRNA properties, we could conclude that no single tRNA property (such as specific wobble pair or modification) was responsible for high readthrough. Nevertheless, we could not rule out synergistic effects of both nucleotide and tRNA on readthrough efficiency. The same arguments applied to our discovery of readthrough-inhibiting candidates: CGU, GCU, UUA, and UUC. Further experiments are needed to deconstruct the roles of nucleotide and tRNA in mediating readthrough efficiency.

Although ribosome recycling was not the focus of our study, our use of profiling data from recycling factor-depleted cells [42] allowed us to observe that some nucleotides and codons in the P site have specific effects in *rli1-d* cells but not in wild-type or eRF1 mutant cells (Figures 5 and 6). Since wild-type and eRF1 mutant cells were both wild-type for recycling factors, these differences may be attributed to the recycling step. As failure to recycle leads to ribosomes re-initiating randomly within <10 nt downstream of the stop codon [42], there is ∼33% chance that re-initiation stays in-frame with the ORF, and there was no way for us to distinguish these footprints from actual readthrough footprints. Thus, we speculated that the differences in “readthrough” efficiency between recycling factor-depleted cells and other samples could mean these nucleotides or codons affect the recycling step. If this is true, cautious use and interpretation of dual/bi-cistronic reporter assays in measuring readthrough efficiency may need to be exercised, as re-initiation that is in-frame with the original reading frame could still produce a functional readthrough reporter. This is evidenced by previous claims borne out of dual reporter experiments stating that functions of Rli1 and Hcr1 were to control termination and readthrough [53, 54], while ribosome profiling data and subsequent experiments demonstrated their ribosome recycling properties [42, 55]. Unless the readthrough product was defined by the combined size of both internal control and readthrough reporters, enzymatic or colorimetric assays that separately detect the internal control reporter and readthrough reporter may not be ideal methods for studying readthrough contexts.

Another mRNA feature that we found to be a significant determinant of readthrough efficiency was 3’-UTR length. Intriguingly, readthrough efficiency decreased with 3’-UTR length in WT cells but increased with 3’-UTR length in eRF1 mutant cells. The trend in eRF1 mutant cells is consistent with the hypothesis that proximity of a stop codon to PABP enhances termination efficiency [39]. The fact that only eRF1 mutant cells displayed such a trend implied that PABP’s expected role in enhancing termination is principally observable when eRF1 function is not efficient. This result is consistent with recent work from our laboratory demonstrating that deletion of the yeast *PAB1* gene led to a reduced accumulation of premature termination products, increased readthrough at multiple PTCs, and increased readthrough of an otherwise “weak” PTC in parallel with 3’-UTR lengthening of the respective transcripts [39]. We surmise that the proximity of the stop codon to PABP, which is thought to enhance termination efficiency through interaction of PABP with eRF3 [37, 56], is masked in the ribosome profiling data of WT cells where eRF1 is fully functional, but when this protein became less efficient or absent, PABP’s role in recruiting eRF3 or other factors could be distinguished. The relationship between PABP’s proximity to a stop codon and the efficiency of termination parallels that of PABP and mRNA stability in the “faux 3’-UTR” model for NMD, where increasing PABP’s distance from a stop codon led to aberrant termination and destabilized the mRNAs [43, 57].

Although normal and premature termination processes have been shown to differ in efficiency, they share the fundamental requirement for release factors and a stop codon [39, 43, 57]. While our study largely involved normal termination codons, we provided additional features of the mRNA sequence that influence termination and readthrough efficiencies, which could be useful in designing or understanding therapeutic approaches for the large number of diseases caused by nonsense mutations [4].

## MATERIALS AND METHODS

### Yeast strains and culture growth conditions

Yeast strains used in this study are in the W303 background. Wild-type (HFY114) and temperature-sensitive *sup45-2* (HFY1218) cells were grown in 1 L of YPD at 25°C with shaking. When the OD600 of the culture reached 0.6-0.8, its temperature was shifted as follows: Cells were collected by centrifugation at 5,000 rpm for 5 min at room temperature (RT) using a JA10 rotor in a Beckman Coulter Avanti J-E centrifuge, resuspended in 400 ml of fresh YPD media, transferred to a new flask, and incubated in a 25°C water bath with shaking. After 20 min of incubation, 400 ml of pre-warmed (57°C) YPD were added to the flask, and cells were incubated in a 37°C water bath for 30 minutes with shaking. As a control for the temperature shift procedure, the same protocol was carried out except that YPD was pre-warmed to 25°C and the subsequent 30-minute incubation was at 25°C.

### Library preparation and sequencing

Ribosome profiling and RNA-seq libraries were prepared as described previously in our library preparation protocol for immunoprecipitated ribosomes [58]. Briefly, 1 L of cells of OD600 between 0.6 and 0.8 were harvested by rapid vacuum filtration and flash-frozen in liquid nitrogen in the presence of Footprinting Buffer (20 mM Tris-HCl pH 7.4, 150 mM NaCl, 5 mM MgCl2) plus 1% TritonX-100, 0.5 mM DTT, 1 mM phenylmethylsulfonyl fluoride (PMSF) and 1X protease inhibitors. Cells were lysed in a Cryomill (Retsch) at 5 Hz for 2 min, then 10 Hz for 15 min. Lysates were clarified by ultracentrifugation in a Beckman Coulter Optima L-90K Ultracentrifuge at 18,000 rpm for 10 min at 4°C, using a 50Ti rotor. Centrifugation was repeated for the supernatant at 18,000 rpm for 15 min at 4°C.

For ribosome profiling [59], lysates were digested with RNaseI (Invitrogen, #AM2294), then layered onto a 1 M sucrose cushion in Footprinting Buffer plus 0.5 mM DTT and centrifuged in a Beckman Optima TLX Ultracentrifuge at 60,000 rpm for 1 hour at 4°C using a TLA100.3 rotor to isolate 80S ribosomes. Ribosome-protected fragments were extracted using a miRNeasy kit (QIAgen, #217004) and depleted of rRNA using a Yeast Ribo-Zero Gold rRNA removal kit (Illumina, #MRZY1324) according to the manufacturer’s protocol. Multiplexed cDNA libraries were prepared with the NEXTFlex Small RNA-Seq Kit v3 (BIOO Scientific, #NOVA-5132-06) according to the manufacturer’s protocol and sequenced (single-end, 75 cycles) on a NextSeq500.

For RNA-Seq, total RNAs were extracted from lysates using a miRNeasy kit (QIAgen, #217004), depleted of genomic DNA contamination using Baseline-Zero DNase (Epicentre, #DB0715K), and depleted of rRNA using a Yeast Ribo-Zero Gold rRNA removal kit (Illumina, #MRZY1324) according to the manufacturer’s protocol. Multiplexed cDNA libraries were prepared with a TruSeq Stranded mRNA sample prep kit (Illumina, #RS-122-2101) according to the manufacturer’s protocol and sequenced (single-end, 75 cycles) on a NextSeq500.

### Data processing

The following software packages were used to process RNA- and ribo-seq library sequences: fastqc v0.10.1 (https://www.bioinformatics.babraham.ac.uk/projects/fastqc/); cutadapt v1.9 [60]; samtools v0.1.19 [61]; bowtie v1.0.0 [62], cufflinks v2.2.1 [63]; bam2fastq v1.1.0 (https://gsl.hudsonalpha.org/information/software/bam2fastq); RSEM v1.3.0 [64].

A customized yeast transcriptome was constructed from the S288C reference genome sequence and annotations downloaded from the Saccharomyces Genome Database (https://www.yeastgenome.org) on September 10, 2015 (S288C_reference_genome_R64-2-1_20150113). Annotations for the following classes of transcripts were parsed from the genomic gff3-formatted annotation file and processed separately: protein coding genes (6551); intron-containing genes (272); genes with 5’-UTR introns (24); frameshifted genes (47); blocked and pseudogenes (18), non-coding RNAs (ncRNAs) (16). For all but the last class, 5’- and 3’-UTR entries were added to the individual gff files employing the UTR annotations from Nagalakshimi et al. [65]. 5’- and 3’-UTR information was available for 2840 and 2849 genes, respectively. In cases where no UTR was annotated, we used the 75^th^ percentile lengths of 97 and 173 nt, respectively. Unspliced versions (pre-mRNAs) of all intron-containing genes were defined as a continuous exon extending from the start of the 5’-UTR to end of the 3’-UTR. After editing the individual parsed gff files, they were concatenated, and a fasta file with sequences of all 6952 transcripts was generated using the cufflinks gffread program. Information included in the fasta header lines generated by gffread was used to construct a transcript-to-gene-map file for use with RSEM, which was used for gene- and isoform-level quantification of transcripts. A modified transcriptome was used for riboWaltz (see below). The riboWaltz transcriptome contained only the spliced transcripts of protein coding genes (6551) and genes with 5’-UTR introns (24). In addition, the riboWaltz transcriptome considered the stop codon as a part of the 3’-UTR region rather than the CDS region, thus the length of CDS and 3’-UTR regions of each gene were adjusted accordingly.

For ribosome profiling libraries, de-multiplexed sequences were first processed by cutadapt to filter out low quality reads and read lengths shorter than 23 nt while trimming adapter sequences (cutadapt -a TGGAATTCTCGGGTGCCAAGG -q 10 --trim-n -m 23). Reads aligning to non-protein coding RNAs (rRNA, tRNA, snoRNA, and other non-protein coding RNA sequences) were removed by aligning with bowtie (-m 100 -3 4 -5 4 -n 2 -l 15 --un) and retaining only unaligned reads. The random 4-mers introduced on both sides of each RPF during library preparation were trimmed, concatenated, and stored as an identifier “barcode” used to remove PCR duplicates by aligning reads to the transcriptome (bowtie -m 10 -n 2 -l 15 -S --best --strata) and retaining only a single barcode (and associated read) at each position where multiple reads aligned. After duplicates removal, the SAM-formatted alignment file was converted either to BAM format using samtools or to fastq format using bam2fastq for use in downstream analyses. Transcript abundance measurements were determined by RSEM with settings --seed-length 15 --bowtie-m 10. Reads processing and alignment statistics can be found in Table S2.

Subsequent analyses of ribosome profiling data were performed in the R software environment (versions 3.5 and 3.6). RPFs of 20-23 nt and 27-32 nt in length were used for the analyses. P-site offsets of reads in ribosome profiling libraries were assigned using the R package riboWaltz [66], the results of which are provided in Table S3. After assigning P-site location, reads from replicate libraries were combined. In addition to riboWaltz, the following packages were used for data analysis and visualizations: data.table, dplyr, reshape2, seqinr, ggplot2, and scales. Built-in base R functions, rcompanion, and ggpubr were used for statistical analyses. randomForest and caret were used to generate and analyze random forest models.

The TruSeq stranded RNA-Seq libraries were processed using RSEM with no additional pre-processing.

### Public ribosome profiling data sets

Public ribosome profiling data of *sup45-d* [41], *rli1-d* [42], and their WT counterparts were downloaded from NCBI’s Gene Expression Omnibus (GEO) database. Yeast strain genotypes, cell growth conditions, accession numbers of the data series and specific sequencing runs are outlined in Table S1. After download, reads were adapter-trimmed and length-filtered by cutadapt (cutadapt -a CTGTAGGCACCATCAAT -q 10 -- trim-n -m 15), after which reads aligning to non-protein coding RNAs were removed (bowtie -m 100 -n 2 -l 15 --un) and remaining reads aligned to the transcriptome exactly as described above, with the omission of the PCR duplicate removal step. Reads processing and alignment statistics can be found in Table S2.

## ACCESSION NUMBERS

The raw sequencing data are deposited and available at Gene Expression Omnibus (GEO) under accession number GSE162780.

## SUPPLEMENTARY DATA

Supplementary data are available at …

## ACKNOWLEDGMENTS

This work was supported by grants to A.J. (5R01 GM27757-37 and 1R35GM122468-04) from the U.S. National Institutes of Health. We thank Jill Moore, Zhiping Weng, Andrei Korostelev, Barry Cooperman, and Sean Ryder for helpful discussions.

## AUTHOR CONTRIBUTIONS

K.M., F.H., R.G., and A.J. conceived and designed the experiments, K.M. carried out the experiments and data analyses, R.B. wrote data processing scripts, K.M. and A.J. wrote the paper with input from all authors, and A.J. obtained funding for the study.

---

**Figure S1.**
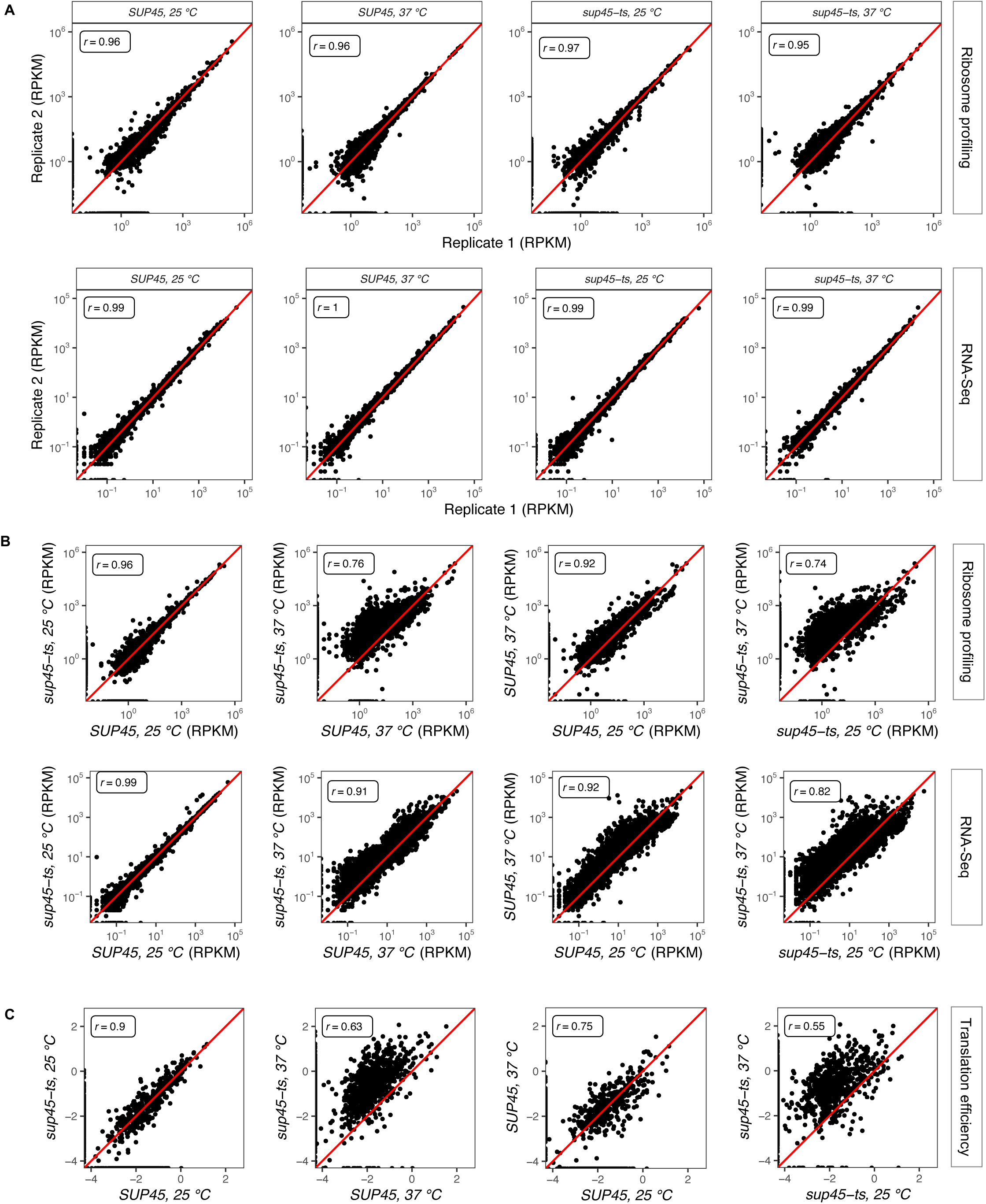
Reproducibility between replicates of *sup45* temperature-shift ribosome profiling and RNA-Seq datasets and changes in global ribosome density, mRNA abundance and translation efficiency upon eRF1 inactivation. **A.** Correlation between ribosome density (top) or mRNA abundance (bottom) in reads per kilobase per million (RPKM) of a pair of replicates for each yeast strain and growth temperature. Pearson’s correlation coefficient (r) was calculated and reported for each pair of replicates. **B.** Correlation between ribosome density (top) or mRNA abundance (bottom) in reads per kilobase per million (RPKM) of different pairs of yeast strains and growth temperatures. Pearson’s correlation coefficient (r) was calculated and reported for each pair of samples. **C.** Correlation between translation efficiency (ribosome density divided by mRNA abundance, log-transformed) of different pairs of yeast strains and growth temperatures. Pearson’s correlation coefficient (r) was calculated and reported for each pair of samples.

**Figure S2.**
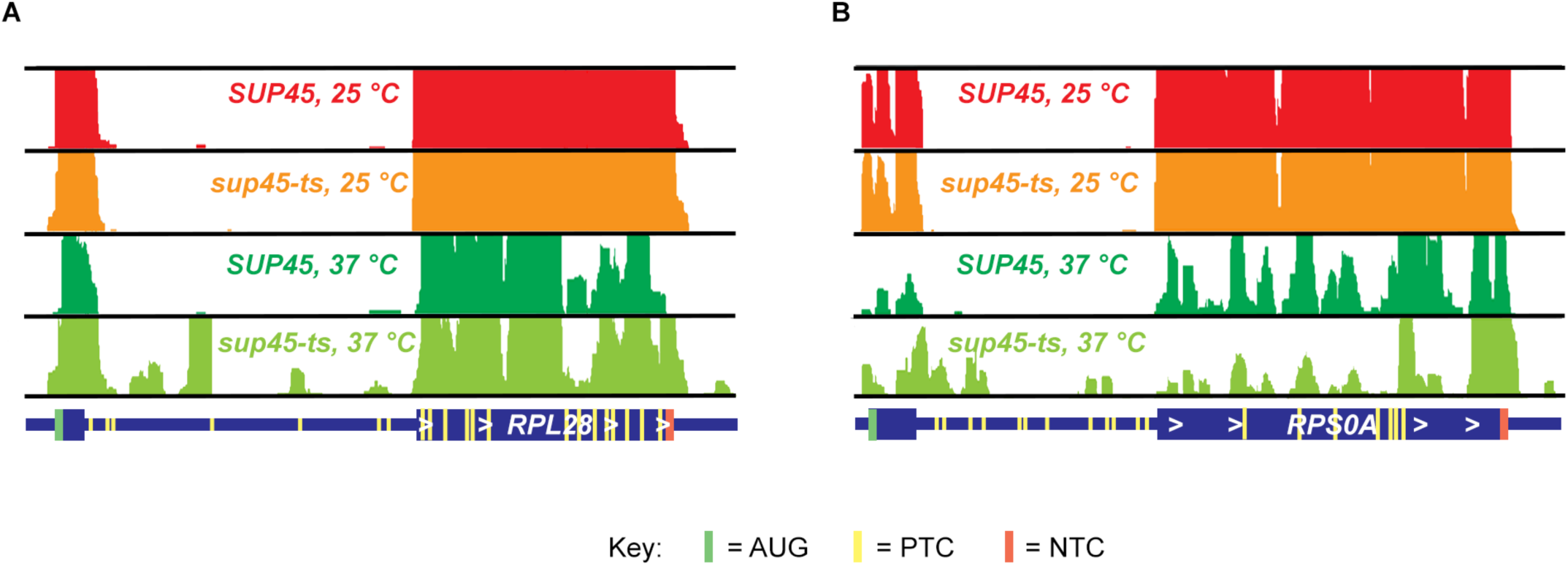
Ribosome occupancy in the transcripts of two intron-containing genes. Read coverage tracks from the Integrative Genomics Viewer (IGV) [67] showing coverage of ribosome profiling reads for the intron-containing genes, **A.** *RPL28* and **B.** *RPS0A,* in *SUP45* and *sup45-ts* strains at 25°C and 37°C. Yellow rectangles indicate the position of termination codons in frame with the respective initiation codons under conditions where the introns are translated. Full scale for A and B equals 50 reads.

**Figure S3.**
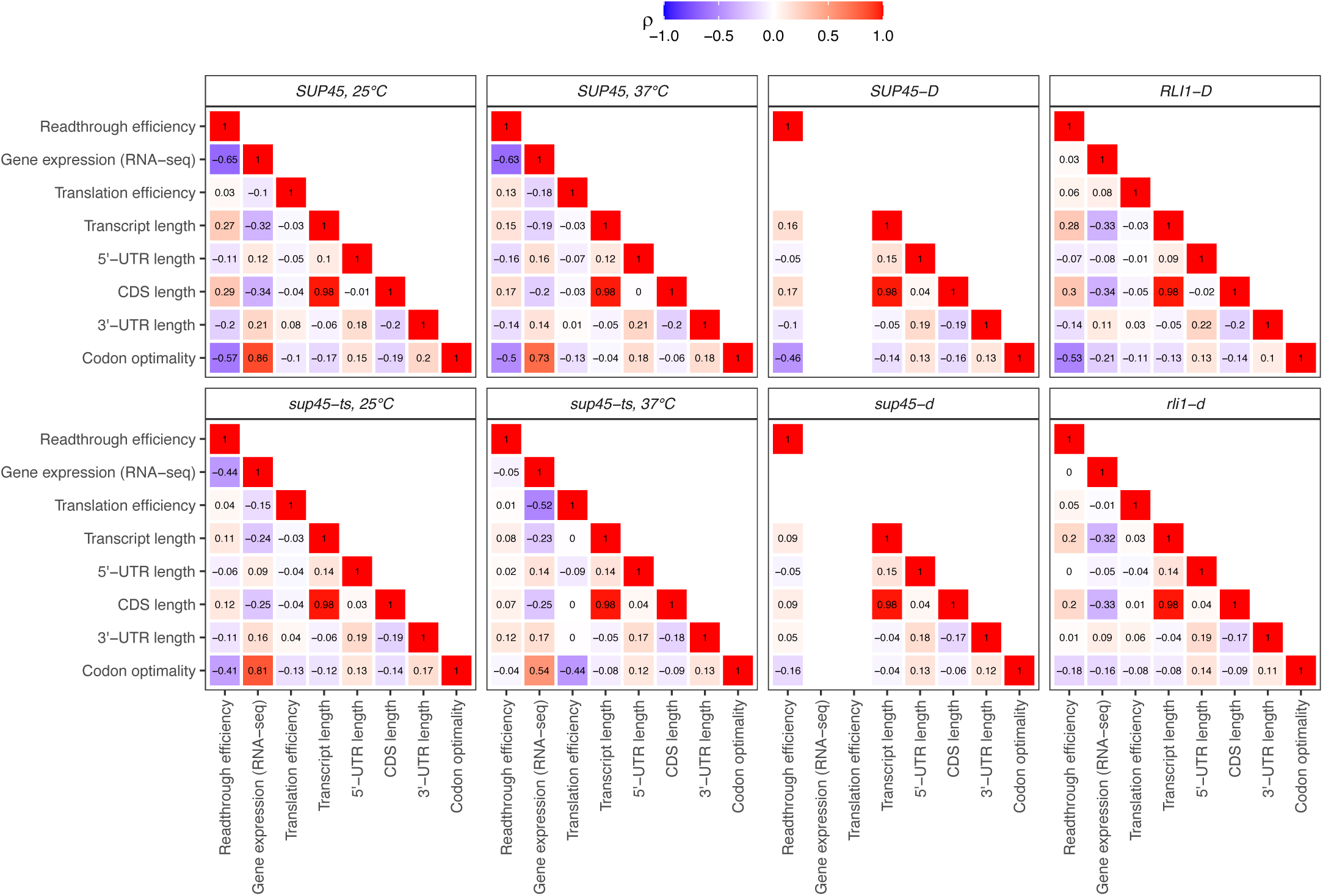
Correlation matrix of relationships between readthrough efficiency, gene expression, translation efficiency, codon optimality, and transcript length. Spearman’s correlation coefficient (ρ) was calculated and reported for each pair of variables.

**Table S1.** Ribosome profiling data and yeast strains.

| Sample | SRR Run | Yeast strain | Notation in this paper | Genotype | Culture media | CHX in lysis buffer |
| --- | --- | --- | --- | --- | --- | --- |
| ***eRF1 temperature-sensitive mutant (This study): GSE162780*** |  |  |  |  |  |  |
| GSM4959785,<br>GSM4959786,<br>GSM4959787,<br>GSM4959788 |  | WT<br>(HFY114) | *SUP45* | *MATa ade2-1 his3-11,15 leu2-3,112 trp1-1 ura3-1 can1-100* | YPD | No |
| GSM4959789,<br>GSM4959790,<br>GSM4959791,<br>GSM4959792 |  | eRF1 ts<br>mutant<br>(HFY1218) | *sup45-ts* | *MATa ade2-1 his3-11,15 leu2-3,112 trp1-1 ura3-1 can1-100 sup45-2* | YPD | No |
| ***eRF1 depletion (Wu et al., 2019): GSE115162*** |  |  |  |  |  |  |
| GSM3168380,<br>GSM3168381 | SRR7241903,<br>SRR7241904 | WT<br>(yCW30) | *SUP45-D* | *MATa his3Δ1 leu2Δ0 met15Δ0 ura3Δ0 HO::ADH1p-OsTIR1-URA3* | YPGR → YPD<br>+ 0.5mM auxin | Yes |
| GSM3168385,<br>GSM3168386 | SRR7241908,<br>SRR7241909 | eRF1<br>depleted<br>(yKW13) | *sup45-d* | *MATa his3Δ1 leu2Δ0 met15Δ0 ura3Δ0 KanMX4:P<sub>GAL</sub>1-SUP45* | YPGR → YPD | Yes |
| ***Rli1 depletion (Young et al., 2015): GSE69414*** |  |  |  |  |  |  |
| GSM1700885 | SRR2046309,<br>SRR2046310 | WT<br>(BY4741) | *RLI1-D* | *MATa his3Δ1 leu2Δ0 met15Δ0 ura3Δ0* | YPG → YPD | Yes |
| GSM1700886,<br>GSM1700891 | SRR2046311,<br>SRR2046312,<br>SRR2046319 | Rli1 depleted<br>(YDH369) | *rli1-d* | *MATa his3Δ1 leu2Δ0 met15Δ0 ura3Δ0 lys2Δ0 rli1Δ::kanMX4 pDH181 (P<sub>GAL</sub>-UBI-R-FH-RLI1, LEU2)* | YPG → YPD | Yes |

**Table S2.** Reads processing and alignment statistics.

| Sample | SRR Run | Total reads processed | Reads ≥ 15 bp & contained adapters | Reads not aligned to ncRNAs* | Duplicate reads | Reads aligned to transcriptome |
| --- | --- | --- | --- | --- | --- | --- |
| *SUP45*, 25°C (replicate 1) |  | 67,532,129 | 57,853,137 | 25,787,398 | 3,441,725 | 20,407,888 |
| *SUP45*, 25°C (replicate 2) |  | 61,556,833 | 51,451,140 | 18,867,321 | 7,016,422 | 10,101,506 |
| *SUP45*, 37°C (replicate 1) |  | 49,805,337 | 44,462,117 | 14,274,514 | 1,866,021 | 11,195,529 |
| *SUP45*, 37°C (replicate 2) |  | 64,613,280 | 55,971,740 | 24,346,626 | 5,660,607 | 16,685,974 |
| *sup45-ts*, 25°C (replicate 1) |  | 91,931,471 | 70,972,671 | 34,574,953 | 5,747,485 | 24,437,810 |
| *sup45-ts*, 25°C (replicate 2) |  | 60,891,476 | 49,432,024 | 25,181,918 | 4,148,599 | 18,939,808 |
| *sup45-ts*, 37°C (replicate 1) |  | 77,154,540 | 55,550,470 | 21,351,921 | 3,510,551 | 14,378,321 |
| *sup45-ts*, 37°C (replicate 2) |  | 66,416,665 | 48,944,777 | 17,330,578 | 3,701,109 | 10,930,634 |
| *SUP45-D* (replicate 1) | SRR7241903 | 54,157,010 | 53,979,859 | 39,261,094 | N/A | 39,261,094 |
| *SUP45-D* (replicate 2) | SRR7241904 | 32,548,315 | 32,188,657 | 21,139,968 | N/A | 21,139,968 |
| *sup45-d* (replicate 1) | SRR7241908 | 46,687,648 | 45,226,887 | 34,527,078 | N/A | 34,527,078 |
| *sup45-d* (replicate 2) | SRR7241909 | 32,916,353 | 32,750,862 | 20,325,474 | N/A | 20,325,474 |
| *RLI1-D* (replicate 1) | SRR2046309 | 52,021,032 | 51,768,936 | 17,545,223 | N/A | 17,545,223 |
| *RLI1-D* (replicate 2) | SRR2046310 | 52,026,256 | 49,283,307 | 41,880,568 | N/A | 41,880,568 |
| *rli1-d* (replicate 1) | SRR2046311 | 46,386,847 | 45,694,917 | 15,064,692 | N/A | 15,064,692 |
| *rli1-d* (replicate 2) | SRR2046312 | 158,583,030 | 155,589,606 | 53,259,434 | N/A | 53,259,434 |
| *rli1-d* (replicate 3) | SRR2046319 | 35,129,965 | 33,633,653 | 28,658,467 | N/A | 28,658,467 |

\*ncRNAs refer to non protein-coding RNAs

**Table S3.** Offset from 5’ and 3’ ends of footprint to the first nucleotide in the P-site for each footprint length in each sample.

| Sample | P-site Offset from 5'/3' ends for each footprint length (nt) |  |  |  |  |  |  |  |  |  |
| --- | --- | --- | --- | --- | --- | --- | --- | --- | --- | --- |
|  | 20 | 21 | 22 | 23 | 27 | 28 | 29 | 30 | 31 | 32 |
| *SUP45*, 25°C (replicate 1) | 12/7 | 13/7 | 13/8 | 14/8 | 11/15 | 12/15 | 13/15 | 13/16 | 13/17 | 13/18 |
| *SUP45*, 25°C (replicate 2) | 12/7 | 13/7 | 13/8 | 14/8 | 11/15 | 12/15 | 13/15 | 13/16 | 13/17 | 13/18 |
| *SUP45*, 37°C (replicate 1) | 12/7 | 13/7 | 13/8 | 13/9 | 11/15 | 12/15 | 13/15 | 13/16 | 13/17 | 13/18 |
| *SUP45*, 37°C (replicate 2) | 12/7 | 13/7 | 13/8 | 14/8 | 11/15 | 12/15 | 13/15 | 13/16 | 14/16 | 13/18 |
| *sup45-ts*, 25°C (replicate 1) | 12/7 | 12/8 | 13/8 | 14/8 | 11/15 | 12/15 | 13/15 | 13/16 | 13/17 | 13/18 |
| *sup45-ts*, 25°C (replicate 2) | 12/7 | 13/7 | 13/8 | 14/8 | 11/15 | 12/15 | 13/15 | 13/16 | 13/17 | 13/18 |
| *sup45-ts*, 37°C (replicate 1) | 12/7 | 12/8 | 13/8 | 14/8 | 11/15 | 12/15 | 13/15 | 13/16 | 13/17 | 13/18 |
| *sup45-ts*, 37°C (replicate 2) | 12/7 | 13/7 | 13/8 | 14/8 | 11/15 | 12/15 | 13/15 | 13/16 | 13/17 | 13/18 |
| *SUP45-D* (replicate 1) | 13/6 | 14/6 | 15/6 | 16/6 | 11/15 | 12/15 | 13/15 | 14/15 | 14/16 | 14/17 |
| *SUP45-D* (replicate 2) | 13/6 | 14/6 | 15/6 | 16/6 | 11/15 | 12/15 | 13/15 | 14/15 | 15/15 | 16/15 |
| *sup45-d* (replicate 1) | 13/6 | 14/6 | 15/6 | 16/6 | 11/15 | 12/15 | 13/15 | 14/15 | 14/16 | 14/17 |
| *sup45-d* (replicate 2) | 13/6 | 14/6 | 15/6 | 16/6 | 11/15 | 12/15 | 13/15 | 14/15 | 15/15 | 16/15 |
| *RLI1-D* (replicate 1) | 4/15 | 5/15 | 6/15 | 7/15 | 11/15 | 12/15 | 13/15 | 14/15 | 14/16 | 14/17 |
| *RLI1-D* (replicate 2) | 4/15 | 5/15 | 6/15 | 7/15 | 11/15 | 12/15 | 13/15 | 14/15 | 15/15 | 16/15 |
| *rli1-d* (replicate 1) | 4/15 | 5/15 | 6/15 | 7/15 | 11/15 | 12/15 | 13/15 | 14/15 | 14/16 | 14/17 |
| *rli1-d* (replicate 2) | 4/15 | 5/15 | 6/15 | 7/15 | 11/15 | 12/15 | 13/15 | 14/15 | 14/16 | 14/17 |
| *rli1-d* (replicate 3) | 4/15 | 5/15 | 6/15 | 7/15 | 11/15 | 12/15 | 13/15 | 14/15 | 15/15 | 16/15 |

**Table S4.** Number of genes involved in statistical analyses.

| Sample | Regression | Classification |  |  |
| --- | --- | --- | --- | --- |
|  |  | High | Low | Total |
| *SUP45*, 25°C | 829 | 125 | 125 | 250 |
| *SUP45*, 37°C | 565 | 85 | 85 | 170 |
| *sup45-ts*, 25°C | 1347 | 202 | 202 | 404 |
| *sup45-ts*, 37°C | 2484 | 373 | 373 | 746 |
| *SUP45-D* | 975 | 147 | 147 | 294 |
| *sup45-d* | 2146 | 322 | 322 | 644 |
| *RLI1-D* | 937 | 141 | 141 | 282 |
| *rli1-d* | 2007 | 301 | 301 | 602 |

